# Evolutionary dynamics of sex-biased gene expression in a young XY system: Insights from the brown alga genus *Fucus*

**DOI:** 10.1101/2021.08.12.455804

**Authors:** William J. Hatchett, Alexander Jueterbock, Martina Kopp, James A. Coyer, Susana M. Coelho, Galice Hoarau, Agnieszka P. Lipinska

**Author notes:** Corresponding author; Agnieszka P. Lipinska.

## Abstract

The sex-dependent regulation of gene expression is considered to be an underlying cause of often extensive, sexually dimorphic traits between males and females. Although the nature and degree of sex-biased gene expression has been well-documented in several animal and plant systems, far less is known about the commonality, conservation, recruitment mechanisms and evolution of sex-biased genes in more distant eukaryotic groups. Brown algae are of particular interest for empirical studies on the evolution of sex-biased gene expression, as they have been evolving independently from animals and plants for over one billion years. Here we focus on two brown algal dioecious species, *Fucus serratus* and *Fucus vesiculosus,* where male heterogamety (XX/XY) has recently emerged. Using RNA-seq, we study sex-biased gene expression and discuss different evolutionary forces responsible for the evolution of sex-biased genes. We find that both species evolved masculinized transcriptomes, with sex-biased genes allocated mainly to male reproductive tissue, but virtually absent in vegetative tissues. Conserved male-biased genes were enriched in functions related to gamete production, along with sperm competition and include two flagellar proteins under positive selection. In contrast to female-biased genes, which show high turnover rates, male-biased genes reveal remarkable conservation of bias and expression levels between the two species. As observed in other XY systems, male-biased genes also display accelerated rates of coding sequence evolution compared to female-biased or unbiased genes. Our results imply that evolutionary forces affect male and female sex-biased genes differently on structural and regulatory levels. Similar to evolutionarily distant plant and animal lineages, sex-biased gene expression in *Fucus* evolved during the transition to dioecy to resolve intra-locus sexual conflict arising from anisogamy.

## INTRODUCTION

Males and females can display striking differences in morphology, physiology and behavior. Evolution of these sexually dimorphic traits is thought to be rooted in anisogamy and shaped by sex-specific selection (Hedrick and Temeles 1989; Connallon and Knowles 2005; Ellegren and Parsch 2007; Schärer et al. 2012). Ultimately, the sexes are defined by the gamete size they produce (either many small or fewer larger gametes) and sexual selection is predicted to act differently regarding these two distinct reproductive strategies (Kokko and Jennions 2008; Schärer et al. 2012). Due to the disparity of resources and energy invested by males and females in their reproductive cells, it is hypothesized that sexual selection will be stronger in the sex that makes the smaller, more abundant, and relatively ‘cheaper’ to produce gametes, resulting in male-biased sexual dimorphism (Darwin 1871; Bateman 1948; Parker 1979; Schärer et al. 2012; Andersson 2019). Because males and females share most of their genomic sequence, the expression of sexually dimorphic traits rely largely on the regulation of sex-biased gene (SBG) expression (Ellegren and Parsch 2007; Parsch and Ellegren 2013; Grath and Parsch 2016). Sex-biased expression allows males and females to avoid the constraints of a shared genome and to cope with intra-locus conflict arising from sexually antagonistic selection (Rice 1998; Connallon and Knowles 2005; Ellegren and Parsch 2007; Cox and Calsbeek 2009; Parsch and Ellegren 2013).

Sex-biased gene expression has been well documented across a wide number of animal species such as insects (Zha et al. 2009; Perry et al. 2014; Papa et al. 2017), mammals (Yang et al. 2006; Blekhman et al. 2010; Naqvi et al. 2019), birds (Mank et al. 2007; Mank and Ellegren 2009; Harrison et al. 2015), and recently also in plants (Zemp et al. 2016; Darolti et al. 2018; Cossard et al. 2019) and brown algae (Martins et al. 2013; Lipinska et al. 2015; Monteiro et al. 2019; Müller et al. 2021). It has been shown that SBG expression can vary in strength throughout development, can be detected already at juvenile stages (Thoemke et al. 2005; Magnusson et al. 2011; Ingleby et al. 2014; Perry et al. 2014), and can constitute a large proportion of the transcriptome, with up to 90% in extreme cases (Ranz et al. 2003; Ayroles et al. 2009). Genome-wide expression studies have found that sex-biased genes tend to be more abundant in males (known as a masculinised transcriptome), where they show stronger bias and more rapid turnover rates compared to female-biased genes (Parisi et al. 2003; Ranz et al. 2003; Yang et al. 2006; Voolstra et al. 2007; Martins et al. 2013; Parsch and Ellegren 2013; Harrison et al. 2015; Yang et al. 2016). Apart from faster evolution of the gene expression levels, male-biased genes more often experienced higher divergence rates of protein-coding sequences than female biased or unbiased genes. This is at least partially attributed to stronger positive selection (Zhang et al. 2004a; Grath and Parsch 2012; Allen et al. 2018; Cutter 2018, Connallon and Knowles 2005; Ellegren and Parsch 2007; Mank and Ellegren 2009; Parsch and Ellegren 2013). In contrast, female-biased genes often evolve at similar or slower rates compared with unbiased genes possibly due to larger pleiotropic constraints (Ellegren and Parsch 2007; Zhang et al. 2007; Assis et al. 2012). Altogether, these observations suggest that male traits experience stronger sexual selection and sexual conflict arising from anisogamy (Ranz et al. 2003; Connallon and Knowles 2005; Hayward and Gillooly 2011; Janicke et al. 2016). However, our knowledge about the evolution of sex-biased expression is limited, mainly, to the animal species with conspicuous sexual dimorphism, where separate sexes evolved a long time ago.

Here, we study the evolution of sex-biased gene expression in two brown algal species from the order Fucales, which has recently evolved separate sexes (Serrão et al. 1999; Coyer et al. 2006; Heesch et al. 2021). Brown algae are an interesting group to study the evolution of sexual systems and sex-biased expression because they have been evolving independently of organisms such as animals, fungi and plants for over a billion years (Baldauf 2003). The majority of brown algal species engage in a haploid-diploid life cycle where sex is expressed during the haploid gametophyte generation and controlled by haploid sex chromosomes (UV system) (Coelho et al. 2018). In that respect, Fucales are unique among the brown algae as they represent the only group that underwent a recent shift towards a diplontic life history, in which the short-lived male sperm and female egg are the only haploid stages (Coelho et al. 2019). Moreover, the conversion to diploidy imposed a switch to the diploid sex-determination system from the haploid UV (via a hermaphroditic intermediate) in several families of Fucales around 17.5 Mya (Heesch et al. 2021). While the transition to diploid sex determination from the haploid system seems to be irreversible, further transitions towards hermaphroditism within the diploid lineages are still possible and occurred independently in several genera of the Fucaceae (Heesch et al. 2021).

*Fucus* species have a rather simple structure with the vegetative body consisting of a holdfast, a thallus and the fronds. The fronds contain reproductive receptacles which in dioecious species bear either antheridia (producing motile sperm) or oogonia (producing immotile, large eggs) (Serrão et al. 1999; Coyer et al. 2006; Cánovas et al. 2011) (Fig. 1). The eggs produce pheromones which facilitate gamete-gamete recognition by attracting sperm within a very short distance (Müller and Gassmann 1985) and fertilized zygotes usually settle within one to two meters of the parent (Arrontes 1993; Serrão et al. 1997). The different reproductive structures are the only visible sexually dimorphic trait in *Fucus* in the absence of detailed morphometric measures, so that dioicous species are sexed solely by the presence of male or female gametes (Coyer et al. 2002). We focused on the dioecious species of two distinct lineages, *Fucus serratus* (Lineage 1) and *Fucus vesiculosus* (Lineage 2), that dominate the rocky intertidal North Atlantic shoreline. The two lineages evolved around 0.9 to 2.25 Mya and both contain hermaphroditic species, including *Fucus distichus* (Lineage 1) and *Fucus spiralis* (Lineage 2) (Serrão et al. 1999; Coyer et al. 2006; Hoarau et al. 2007). All four species often occur intertwined with one another (Fig. 1A) and molecular studies have shown that hybridization is common, involving dioecious-hermaphrodite species pairs within each lineage, but hybrids of dioecious species are almost never found (Coyer et al. 2002; Wallace et al. 2004; Billard et al. 2005; Coyer et al. 2007; Hoarau et al. 2015). Character mapping analysis suggested dioecy as the most likely ancestral sexual system in the *Fucus* genus, however, the direction of transition between hermaphroditism and separate sexes within Lineage 1 and 2 remains ambiguous (Heesch et al. 2021).

**Figure 1.**
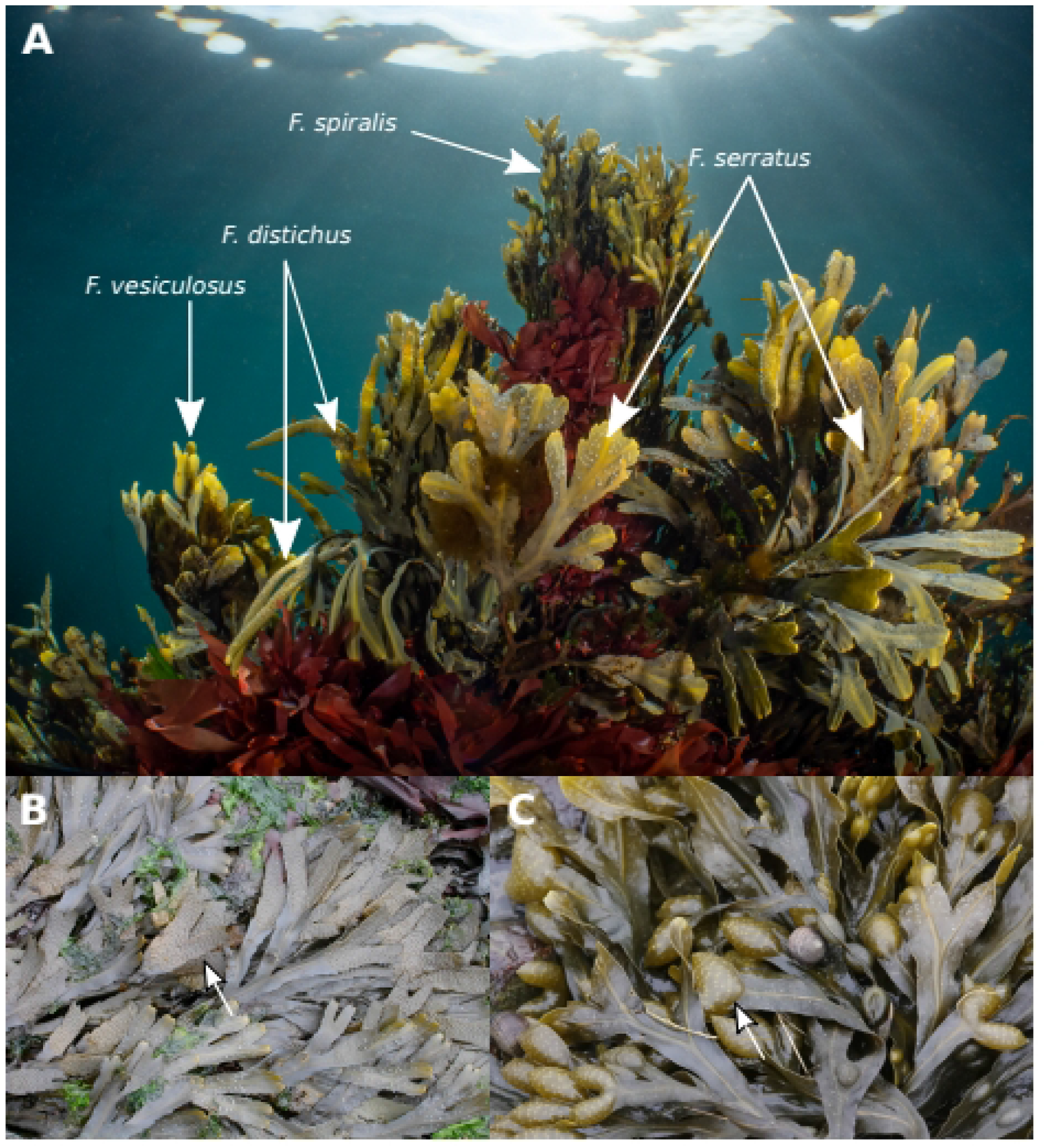
*Fucus* species co-ocurring in their natural habitat. A) *Fucus spiralis* (top), *Fucus distichus* (center-left), *Fucus serratus* (center-right) and *Fucus vesiculosus* (bottom-left) living in sympatry. B) *Fucus serratus* with reproductive structures shown by white arrow. C) *Fucus vesiculosus* with reproductive structures shown by white arrow. Photo credit G. Hoarau (A), J. Coyer (B,C).

Field observations and laboratory crosses of *Fucus serratus–Fucus distichus* hybrids, allowed the identification of the type of sexual system in dioecious species as a male heterogamety (XX/XY) (Coyer et al. 2002). Combined with the low levels of selfing, almost 100% fertilization success in dioecious species and effective polyspermy block (Bolwell et al. 1977; Brawley 1992; Pearson and Brawley 1996; Serrao et al. 1996; Coyer et al. 2002), these observations suggest that the targets of reinforcement and speciation in *Fucus* involve the gamete attraction and/or recognition genes. Moreover, high levels of sperm competition in marine free spawners like *Fucus* imply there is strong selection pressure on the males for reproductive success as species in sympatry have increased sperm specificity (Hoarau et al. 2015).

In this work, we explore male and female transcriptomic data of *Fucus serratus* and *Fucus vesiculosus* which recently evolved dioecy, to elucidate the early stages of the evolution of sex biased gene expression. We study evolutionary dynamics of sex-biased transcriptome expression, investigate the conservation of gene expression patterns between the two algal species and identify sex-biased genes with signatures of positive selection in this relatively young XX/XY system.

## RESULTS

### Transcriptome assembly and analysis of gene expression

We sequenced reproductive and vegetative tissue from males and females of dioecious *F. serratus* and *F. vesiculosus*. We obtained a total of 478 million reads from two sequencing runs with an average of over 21 million reads per tissue type and species (Table S1). The *de novo* assembled reference transcriptome for each species contained 29,610 genes for *F. vesiculosus* and 39,009 genes for *F. serratus* (Table S1, see “Methods” section for details), after filtering out the transcripts with low expression or high similarity to other transcripts. BUSCO v3 (Waterhouse et al. 2018) estimated completeness of each reference transcriptome at 88.8% for *F. vesiculosus* and 92.4% for *F. serratus* (Table S1).

### Sex-biased gene expression

Genes with significant sex-biased expression (FC>=2, p_adj_<0.05) were identified in two comparisons, male reproductive vs female reproductive tissue and male vegetative vs female vegetative tissue, using the DESeq2 R package (Love et al. 2014) (Table S2,S3). As expected, the greater number of sex-biased genes (SBGs) was found in the reproductive tissue when male vs female receptacles were compared (2,993 and 2,772 genes in *F. serratus* and *F. vesiculosus*, respectively) (Fig.2A). In contrast, in vegetative tissues, only 20 and 22 genes were found sex-biased in *F. serratus* and *F. vesiculosus*, respectively (Table S2,S3). Since the sex-biased genes from the vegetative tissue overlapped largely with those from the reproductive tissue, we decided to focus on the latter in all consecutive analyses on sex-biased gene expression.

**Figure 2.**
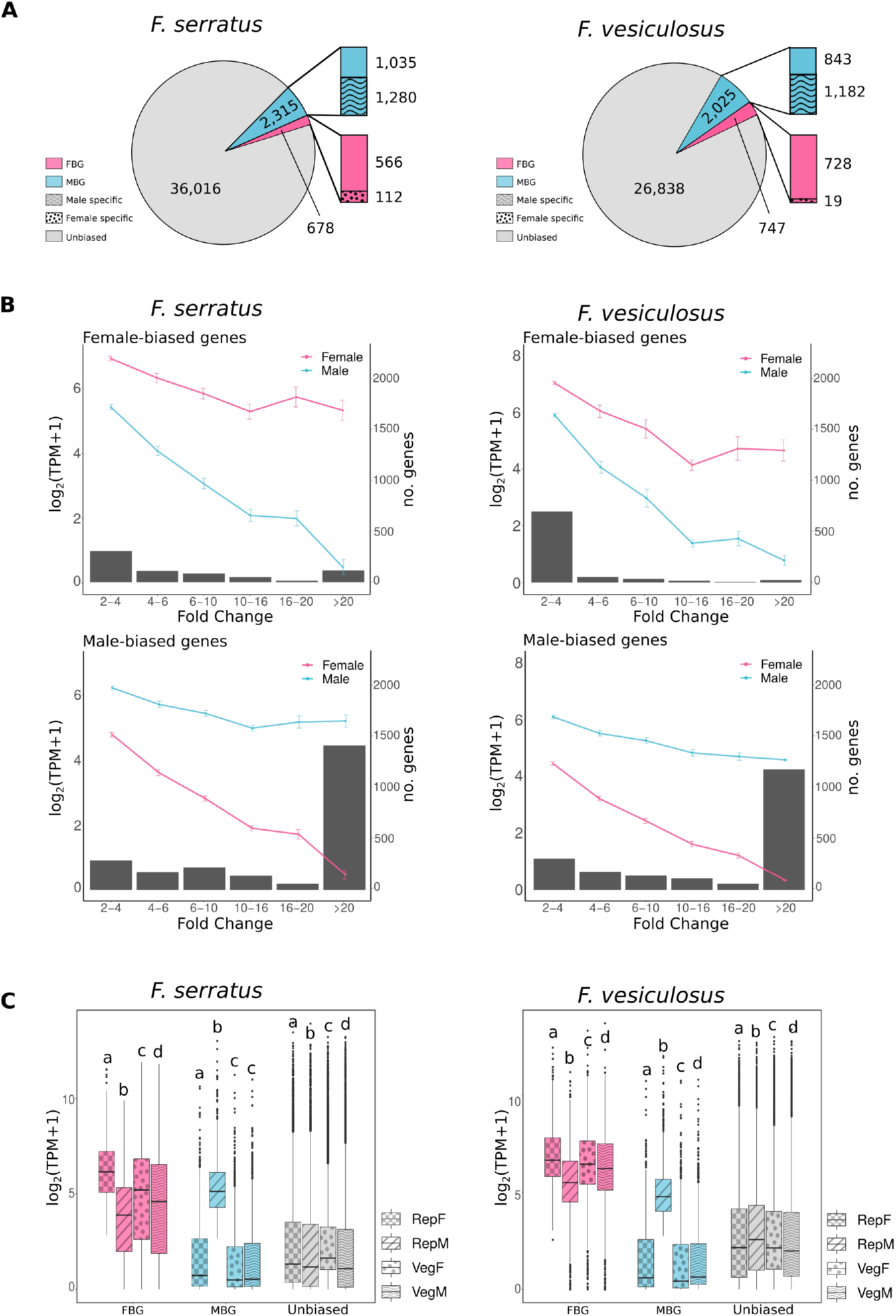
Sex-biased gene expression. A) Number of sex-biased genes (MBG - male-biased and FBG – female-biased) in *F. serratus* and *F. vesiculosus* reference transcriptomes. Unbiased genes were defined as p_adj_>0.05 or showing less than 2-fold difference between the sexes. Bars represent the proportion of sex-specific genes among the sex-biased genes in each species. B) Mean expression levels (log_2_(TPM+1)) of female-biased and male-biased genes at several degrees of sex-bias (Fold Change) in the female (pink) and male (blue) reproductive tissues. Error bars represent standard errors. Bar plot indicates the number of genes in each FC category. *C*) Boxplot showing the mean expression levels across the replicates (log_2_(TPM+1)) of female-biased (pink), male-biased (blue) and unbiased (grey) genes in male and female reproductive and vegetative tissues. The letters above the plots indicate significant differences within each gene group (pairwise Wilcoxon test, p<0.05). RepF - female reproductive tissue; RepM - male reproductive tissue; VegF – female vegetative tissue; VegM – male vegetative tissue.

We found more male-biased genes (MBGs) than female-biased genes (FBGs) in both species (2,315 MBGs vs 678 FBGs in *F. serratus;* and 2,025 MBGs vs 747 FBGs in *F. vesiculosus*) (Fig.2A). Noteworthy, more than half of the MBGs were also male-specific (55% in *F. serratus* and 58% in *F. vesiculosus*), meaning their expression in female reproductive tissue fell below the detection threshold (log_2_(TPM+1)<0) (Fig.2A). In contrast, the majority of female-biased genes were also expressed in male receptacles, and female-specific genes constituted a smaller fraction of the female sex-biased gene (FBG) pool (17%, *F. serratus*; 3%, *F. vesiculosus*) (Fig.2A) (Table S3).

To further examine the relationship between the expression levels and the degree of sex-bias, we grouped the genes according to the Fold Change (FC) difference between males and females and plotted their mean expression levels in each sex (Fig.2B). In both *Fucus* species, sex-biased expression resulted from down-regulation of gene expression in the opposite sex rather than up-regulation in the given sex (Fig. 2B). In the genes showing strongest sex-biased expression (FC>20), down-regulation resulted from silencing (log_2_(TPM+1)<0) of the given gene in the other sex. Interestingly, between 60-90% of female-biased genes featured moderate expression bias (2<FC<6) (416 in *F. serratus*; and 674 in *F. vesiculosus*), whereas the majority of male-biased genes were silent in females and exhibited very high fold changes (FC>20) (61% or 1,416 genes in *F. serratus* and 59% or 1,201 genes in *F. vesiculosus*) which confers with the high proportion of male-specific SBGs (Fig.2A,B).

We also noted that female-biased genes were highly expressed and ubiquitously present in both sexes and both tissue types, including male receptacles (Fig.2C). Conversely, MBGs showed a strong signal of expression only in the male reproductive tissue, and had significantly lower expression levels compared to unbiased genes in male and female vegetative and female reproductive tissues in both species (Fig.2C, p<2e-16 in all pairwise Wilcoxon tests).

### Tissue-biased gene expression

We analyzed transcript abundance in the reproductive versus vegetative tissues within each sex and species to identify genes with tissue-biased expression (FC>=2, p_adj_<0.05) (Table S2,S3)(Fig.3A). Males of both *Fucus* species displayed higher tissue-bias than females, and more of these tissue-biased genes were over-expressed in the reproductive organs compared to vegetative tissue (Fig.3A). To identify sex-biased genes that were predominantly expressed in the reproductive tissue, we compared the tissue-biased data set with that of the male and female sex-biased genes identified above. Not surprisingly, most of the male reproductive tissue biased genes overlapped with MBGs (72% and 88% in *F. serratus* and *F. vesiculosus*, respectively), whereas FBGs were more uniformly expressed across the female body (only 18% and 7% localized specifically in the reproductive tissue of *F. serratus* and *F. vesiculosus,* respectively) (Fig.3A, shaded area). Noteworthy, the SBGs showed significantly higher degrees of sex-bias in the reproductive tissue than in the non-reproductive tissue in both sexes and species (Fig.3B, Wilcoxon test, p<1.4e-06).

**Figure 3.**
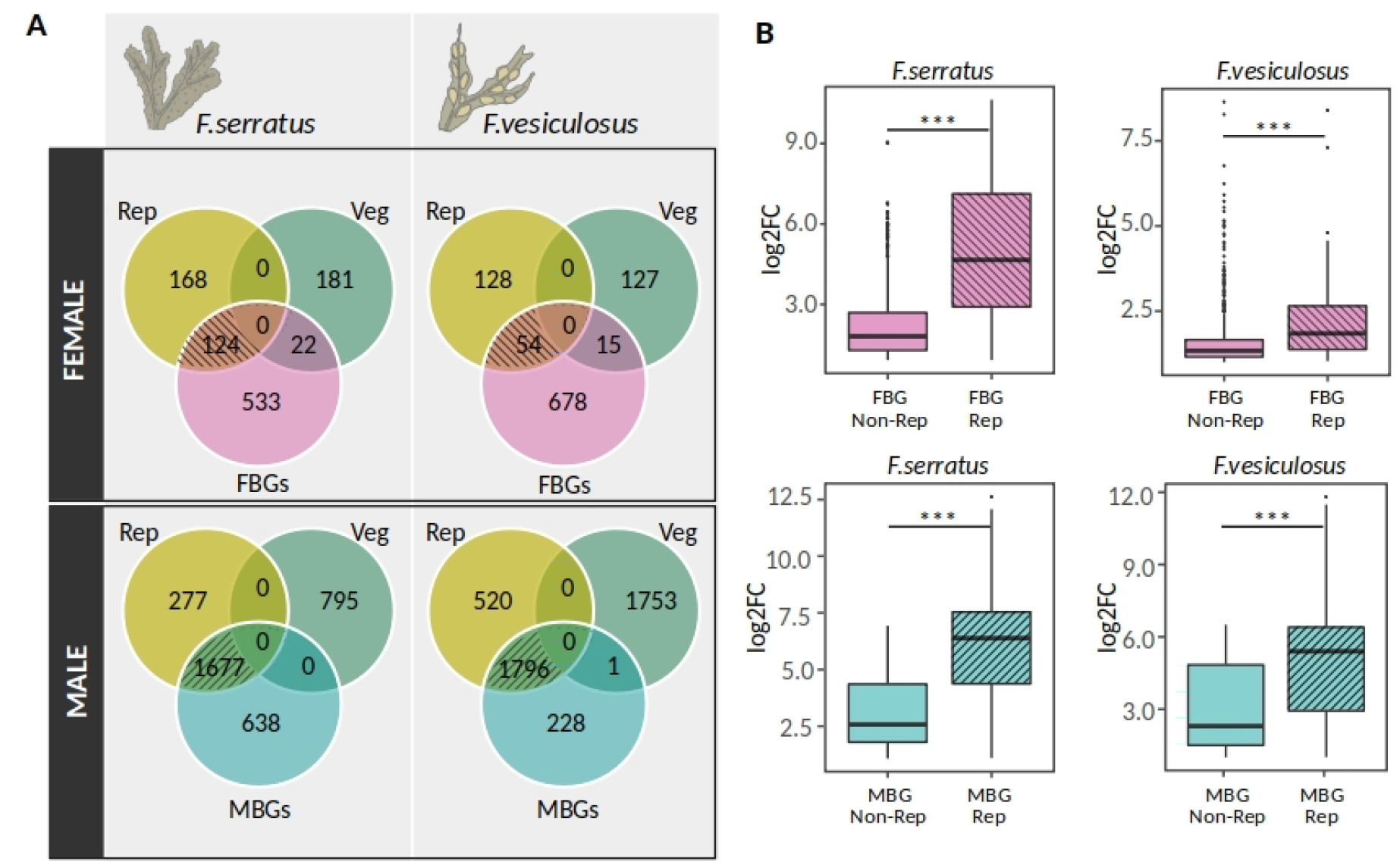
Sex-biased genes (SBGs) are over-expressed specifically in the reproductive tissue. A) Venn diagram shows numbers of significantly differentially expressed genes between reproductive (Rep) and vegetative (Veg) tissues of males and females from *F. serratus* and *F. vesiculosus* (FC>2, p_adj_<0.05). The shaded overlap highlights female-biased genes (top) and male-biased genes (bottom) that were over-expressed in reproductive tissue. B) Overall levels of sex-biased expression (log_2_FC) of SBGs up-regulated in reproductive (Rep) or vegetative tissue (Non-Rep)(̈́(Wilcoxon test, p<1.4e-06).

### High conservation of male-biased expression in *Fucus*

Using Orthofinder, we found 20,077 orthogroups that comprised 85,430 genes (72.6% of all the genes), out of which 14,818 OGs contained genes from both dioecious species (*F. vesiculosus* and *F. serratus)*. In addition, we searched for single copy orthologs within each lineage (*F.distichus* – *F. serratus,* hereinafter Lineage 1, and *F.spiralis* – *F. vesiculosus,* hereinafter Lineage 2) as well as between the two dioecious species (*F. serratus – F. vesiculosus). We* found 9,401 and 8,758 one-to-one orthologs in Lineage 1 and Lineage 2 respectively, and 9,778 one-to-one orthologs in the dioecious pair (Table S4). Up to 35% of genes in each species were “orphans”, meaning species-specific genes, without any intra- or inter-specific orthologs.

Comparisons of orthogroups comprising the sex-biased genes of *F. serratus* and *F. vesiculosus* revealed that the male biased genes were highly conserved between the two species (Table S4). As much as 65% to 75% of the OGs containing male-biased genes were common between *F. serratus* and *F. vesiculosus*. In contrast, only 20% to 26% of orthogroups with female-biased genes were shared between these species (Fig.4A). Interestingly, the low number of conserved female-biased genes was not caused by the presence of orphan genes among FBGs, but rather gain/loss of female bias in existing, orthologous genes. In fact, the proportions of sex-biased genes among the orphan genes were significantly lower than expected in both species and sexes (Chi-square test, p<2.4e-23, Table S5). Taken together, we observed high conservation of male sex-biased expression and higher turnover of female-biased genes between *F. serratus* and *F. vesiculosus*.

**Figure 4.**
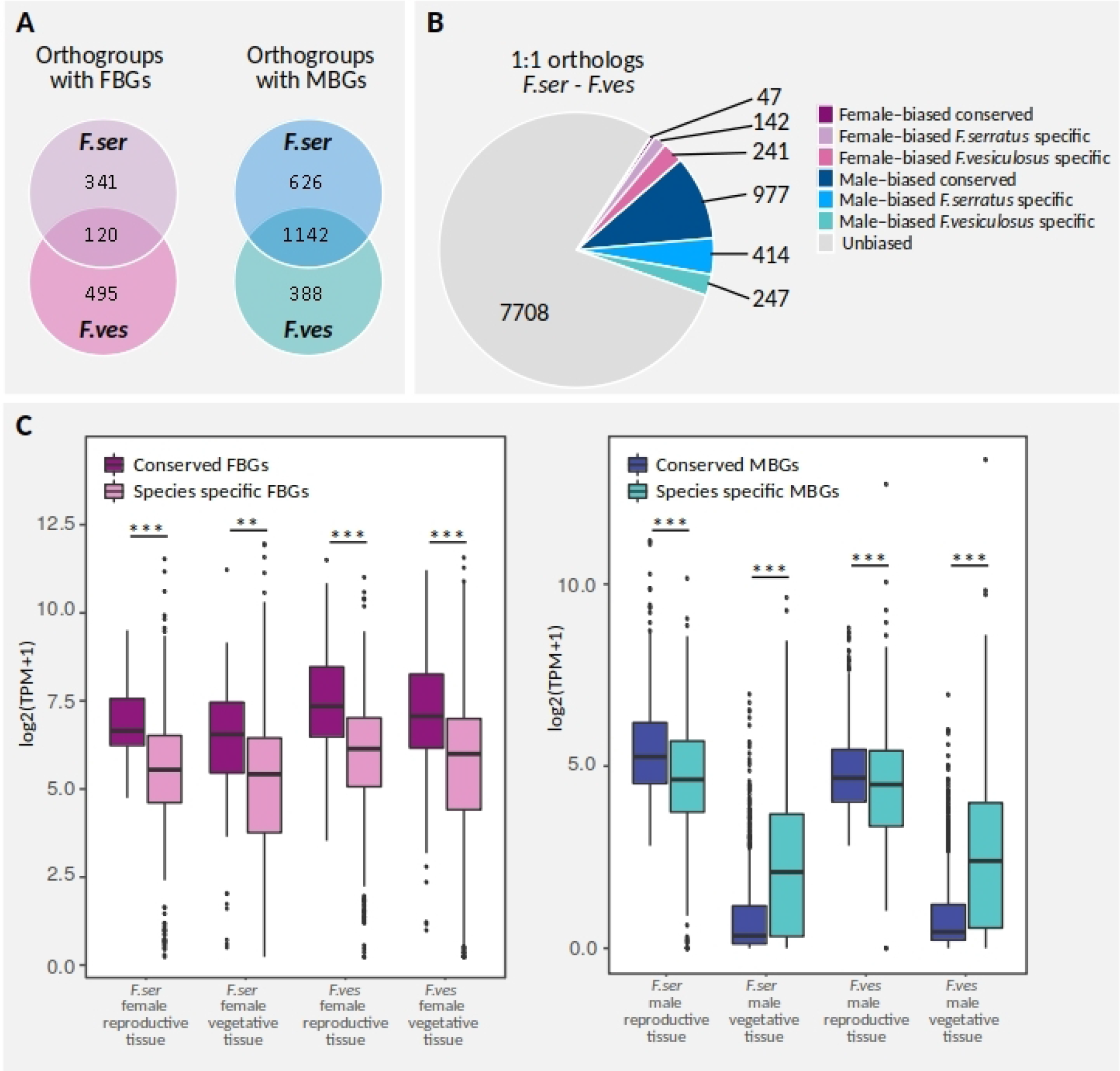
Conservation of sex-biased gene expression across *F. serratus* and *F. vesiculosus* species. A) Numbers of orthogroups with female (pink, FBS) and male (blue, MBS) sex-biased genes shared between dioecious species. Orthogroups with multi-copy genes of a species were included if at least one of the paralogs exhibits sex-biased expression. B) Conservation of sex-biased expression among single copy, one-to-one orthologs between *F. serratus* and *F. vesiculosus*. C) Mean expression levels (log_2_(TPM+1)) of conserved and species-specific SBGs with single copy orthologs in *F. serratus* and *F. vesiculosus* across different tissue types. Wilcoxon test, **p<0.01, ***p<0.001. F.ser – *Fucus serratus*, F.ves – *Fucus vesiculosus*

To further analyze the conservation of the sex-biased expression, we focused on genes for which there was a clear one-to-one relationship across *F. serratus* and *F. vesiculosus*. Out of the 9,778 orthogroups with single copy genes, 21% (2,070 OGs) contained genes with sex-biased expression in at least one of the two species (Table S4). Again, male sex bias was strongly conserved across the two lineages and applied to roughly 70-80% of MBGs with one-to-one orthologs, contrary to 25-16% of conserved FBGs in *F. serratus*/*F. vesiculosus* (Fig.4B).

The patterns of expression of conserved and species-specific SBGs showed similar trends in *F. serratus* and *F. vesiculosus* (Fig.4C). Genes with conserved sex bias had significantly higher average expression levels in reproductive tissue than the species specific SBGs (genes biased towards one sex in one species but not the other) (Fig.4C, Wilcoxon test, p<0.001). Interestingly, this was also true for the conserved FBGs in the vegetative tissue (Wilcoxon test, p<0.01), whereas conserved male biased genes exhibited significantly lower expression levels in the vegetative tissue compared to species-specific MBGs (Wilcoxon test, *p*<0.001). In short, conserved male-biased genes were primarily expressed in reproductive tissue and constituted almost half of the male biased genes found in the receptacles (42% in *F. serratus* and 48% in *F. vesiculosus*).

The tissue specificity of male-biased genes was further highlighted in the hierarchical clustering of the one-to-one orthologs based on expression levels within and among the *F. serratus and F. vesiculosus* species (Fig.5). For the sex-biased genes (at least one or both orthologs are SBGs), the male reproductive samples formed a separate cluster from all the other samples (Fig.5A), which grouped primarily by phylogenetic relatedness, with female reproductive tissue appearing more similar to that of male and female vegetative tissue (Fig.5A). For unbiased genes (neither of the orthologs showed sex-bias) the samples clustered by phylogeny and tissue types (Fig.5B).

**Figure 5.**
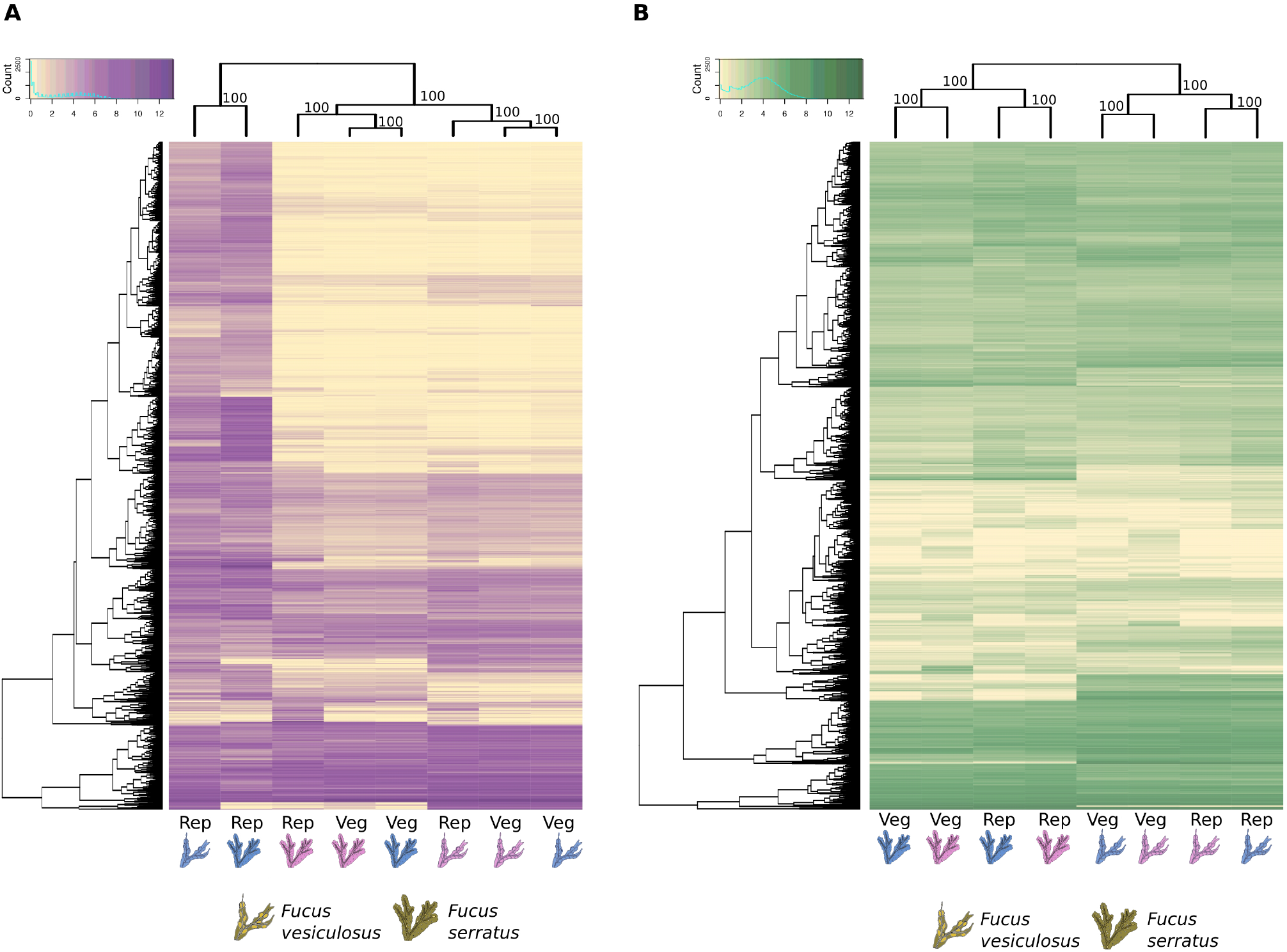
Heatmaps and hierarchical clustering of gene expression levels (log_2_(TPM+1)) for all single copy orthologs among *F. serratus* and *F. vesiculosus*. The dendrogram was generated using hierarchical clustering with 1000 bootstraps (pvclust package, R). A) Sex-biased genes (at least one sex-biased gene in one of the studied species); B) unbiased genes (none of the genes was sex-biased). Rep – Reproductive tissue, Veg – Vegetative tissue, Pink -female, Blue – male

### Evolution of sex-biased genes

To investigate the role of selection on coding sequence evolution, we calculated pairwise divergence of the one-to-one orthologs within lineages (Lineage1: *F. serratus* – *F.distichus* (8,208 orthologs); Lineage2: *F. vesiculosus* – *F.spiralis* (7,363 orthologs)) using the CODEML package in PAML4 (Yang 2007).

In both dioecious species, female-biased genes showed similar rates of non-synonymous to synonymous substitutions (dN/dS) to that of unbiased genes (Fig.6A, Wilcoxon test, *p*>0.25). In contrast, the average dN/ dS was significantly higher for male-biased than unbiased genes (Fig.6A, Wilcoxon test, p<8e−05) and did not depend on the magnitude (FC) or conservation (universal vs species-specific) of the sex-biased expression patterns (Table S6, Wilcoxon test, p>0.11). In addition, in *F. vesiculosus* we found a significant difference in dN/dS ratios between male SBGs and female SBGs (Fig.6A, Wilcoxon test p=0.0026). Although *F. serratus* showed a similar trend, the difference was not significant (Fig.6A, Wilcoxon test p=0.074).

**Figure 6.**
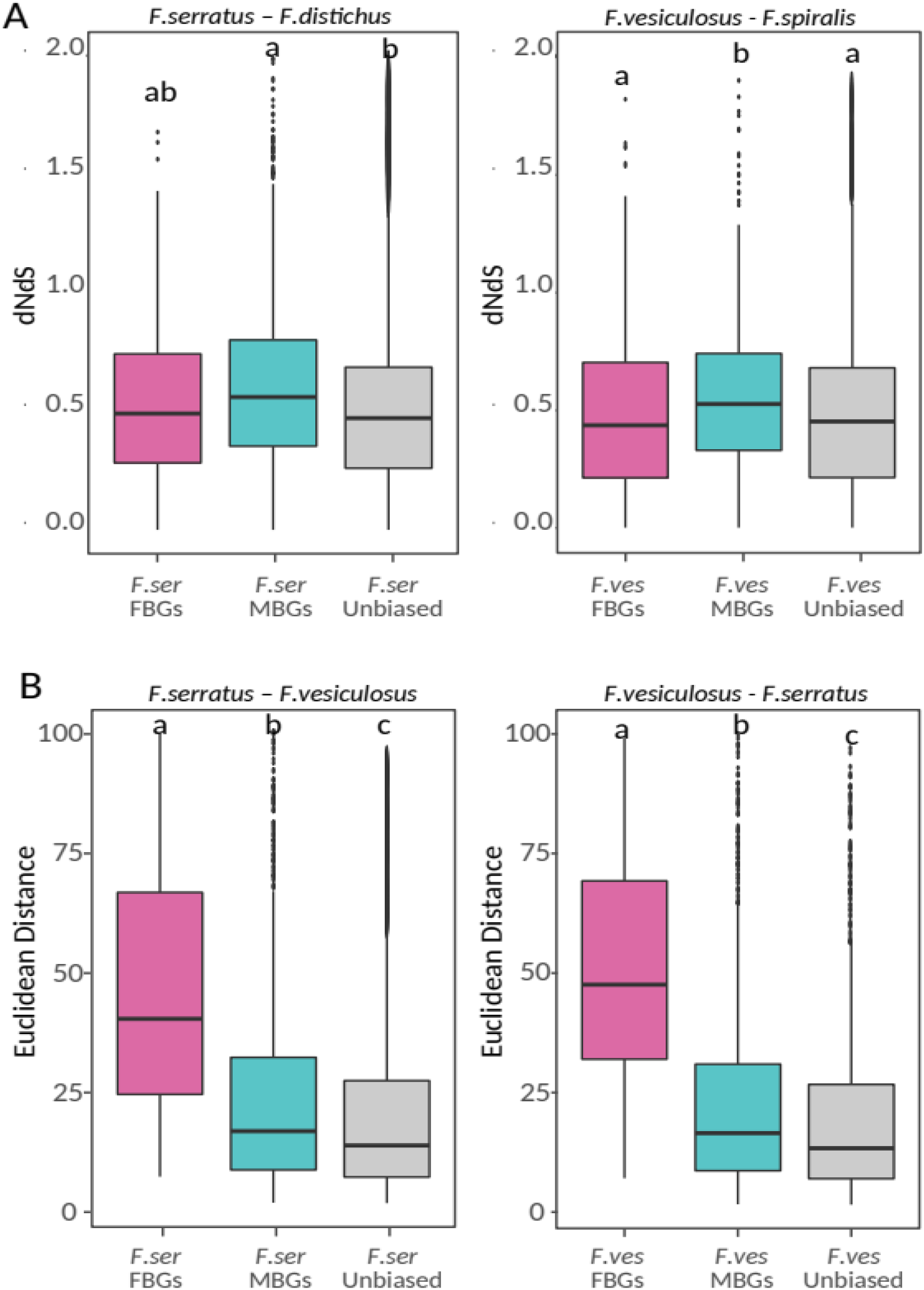
Evolution of sex-biased genes. A) Evolutionary rates measured as dN/dS between species pairs (*F. serratus/F.distichus* and *F. vesiculosus/F.spiralis*) for unbiased, female-biased, and male-biased genes in the two dioecious *Fucus species*. B) Expression divergence measured as Euclidean distances between single copy orthologous genes of *F. serratus* and *F. vesiculosus*. Different letters above the plots indicate significant differences (pairwise Wilcoxon test; p<4.3e-10). FBGs – female-biased genes, MBGs – male-biased genes.

To assess whether increased protein divergence rates were due to increased positive selection or relaxed purifying selection, we performed a maximum likelihood analysis using CODEML in PAML4 **(Yang 2007)**. We used sequences from the four *Fucus* species (*F. vesiculosus*, *F. serratus*, *F. distichus* and *F. spiralis*) and two other brown algae (*Ectocarpus sp*. (Cock et al. 2010) and *Saccharina japonica (*Ye et al. 2015)) to find 561 conserved, single copy orthologs that generated high quality alignments. Among those, 57 orthologs exhibited male-biased expression and 13 exhibited female-biased expression in at least one of the dioecious *Fucus* species (Table S8). Each alignment was tested for the direction and magnitude of selection on amino acid changes using the paired nested site models (M1a, M2a; M7, M8) implemented in PAML4 (CODEML) (Yang 2007). Likelihood ratio test calculated for either one or both pairs of models (M1a–M2a, M7–M8) detected evidence for adaptive evolution in 16 male-biased genes, 2 female-biased genes and 189 unbiased genes (Table S8). These results are consistent with the idea that sex-biased genes are subject to positive selection, although we found no significant enrichment of genes under positive selection among the sex-biased genes compared to unbiased genes (Chi square test, p>0.05). Finally, we compared the dN/dS analysis with gene expression divergence measured as Euclidean distances for the one-to-one orthologous pairs between *F. serratus* and *F. vesiculosus*. Female-biased genes showed the highest divergence in expression patterns compared to male-biased or unbiased genes (Fig.6B, Wilcoxon test p<2e-16). These results are in line with the higher turnover of FBGs (a given gene has a female bias in one species but it is unbiased in the other species). By comparison, male-biased genes presented more conserved expression, with the universal MBGs having overall the most stable expression patterns among all SBGs (Fig.S1, Fig.S2 Wilcoxon test p<0.003).

### Functional analysis

We tested for enrichment of particular molecular functions, biological processes, cellular components, or regulatory pathways in the sex-biased genes of the two *Fucus* species. The Gene Ontologies (GO) associated with female-biased genes in *F. serratus* and *F. vesiculosus* were enriched in biological processes related to cell wall synthesis, translation, transmembrane transport, receptor signaling, photosynthesis, cell homeostasis and establishment of cell polarity (Fisher exact test, p<0.05, Table S7). Interestingly, analysis of male-biased genes of both species identified GO terms related to spermatogenesis and sperm competition in addition to microtubule and flagellar movement categories, as well as photo- and chemotaxis (Fisher exact test, p<0.05, Table S7). Furthermore, two consistently male-biased flagellar associated proteins were found to evolve under positive selection (Table S8). These results are coherent with the reproductive functions of males and females, with MBGs being predominantly involved in male germ cell differentiation, sperm motility and response to pheromones produced by the egg, whereas FBGs being related to the development of a future embryo.

## DISCUSSION

Brown algae are excellent models to study the evolution of sexual systems, as their extraordinary divergence in sex determination mechanisms and sexual dimorphism (ranging from isogamy to oogamy) sets them apart from other eukaryotic groups (Silberfeld et al. 2010; Coelho et al. 2019). In this work, we asked whether the recent emergence of separate sexes, and the transition to a diploid life history in *Fucus serratus* and *Fucus vesiculosus* involved concordant changes in gene expression between males and females. We investigated the proportion of the transcriptome that evolved sex-biased expression in this relatively young XX/XY system with modest sexual dimorphism. We also examined if the evolutionary patterns of sex-biased genes in *Fucus* are convergent with the ones found in well-established XY or ZW systems.

### Sex-biased expression in dioecious *Fucus* species

While very few genes were differentially expressed between male and female vegetative tissue, thousands of genes (ca. 8-9% of *F. serratus* and *F. vesiculosus* transcriptomes, respectively) were differentially expressed in the reproductive tissues. A similar fraction of the genome displayed tissue-biased expression between receptacles and the rest of the body within each sex, allocating the majority of tissue- and sex-biased expression to the reproductive organs. These findings agree with the general trend found in animals and plants where reproductive tissues show the highest expression divergence between sexes (animals: Yang et al. 2006; Yang 2007; Pointer et al. 2013; Harrison et al. 2015; Allen et al. 2018; plants: Song et al. 2017; Darolti et al. 2018). This could be expected in *Fucus,* as sexes are morphologically identical except for their receptacles. The overall moderate levels of SBG expression in *Fucus* (8-9%, compared to > 50% in *Drosophila* (Ranz et al. 2003) and wild turkey (Pointer et al. 2013) may be explained by the low levels of sexual dimorphism, external fertilization mode and, accordingly, more narrow range of sexual selection in both *F. serratus* and *F. vesiculosus* (Luthringer et al. 2014). In birds, the proportion of SBGs corresponded with the strength of selection and the extent of phenotypic dimorphism between males and females (Harrison et al. 2015). Similarly, in a male feminized mutant strain of the brown alga *Macrocystis*, sex-specific phenotypes (male, female or feminized male variant) showed sex-specific transcriptomic patterns (Müller et al. 2021).

It is worth noting that the proportion of SBGs in the oogamous *Fucus* much exceeded that of the near-isogamous *Ectocarpus* (Lipinska et al. 2015). *Ectocarpus* is a filamentous brown alga with low levels of sexual dimorphism between the male and female gametophytes, has a haploid–diploid life cycle and produces morphologically similar, small, flagellated male and female gametes (Luthringer et al. 2014; Lipinska et al. 2015). In *Ectocarpus,* obscure phenotypic sexual dimorphism corresponded to less than 4% (658) of genes being sex-biased during the reproductive stage compared to 8% (2,993) in *F. serratus* and 9% (2,772) in *F. vesiculosus* in our study. Furthermore, in oogamous kelp *Macrocystis*, where male and female gametophytes have visibly distinct morphologies (Müller et al. 1979), sex-biased gene expression analysis found 24% (5,442) of genes with male/female bias (Müller et al. 2021). In summary, our results suggest that the evolution of anisogamy alone, without the other morphologically dimorphic characters, has triggered a significant increase in sex-biased gene expression.

### Masculinization of the *Fucus* transcriptome

In both systems, *Ectocarpus* with UV, and *Fucus* with XX/XY sex chromosomes, we identified an excess of male-biased over female-biased genes. However, in *Fucus* species male overexpression was much more pronounced, exceeding more than three times the number of FBGs (400 MBGs vs 258 FBGs in *Ectocarpus*; 2,315 MBGs vs 678 FBGs in *F. serratus*; 2,025 MBGs vs 747 FBGs in *F. vesiculosu*s). Globally, male-biased genes featured extreme expression bias (FC>20) with more than half of the male-biased genes being male specific, expressed explicitly in male receptacles, and at significantly lower levels than unbiased genes in the vegetative tissue. This masculinization of the transcription profile may result from adaptive changes in males, and, as predicted for anisogamy, implies that males experience stronger selection than females (Darwin 1871; Bateman 1948; Parker 1979; Schärer et al. 2012; Andersson 2019). Masculinization of the transcriptome has been found in many other species and could be due to the relative expression of male sexual traits, female choice and male-male competition (Connallon and Knowles 2005; Pointer et al. 2013; Harkess et al. 2015; Zemp et al. 2016). Although female choice in the ‘classical’ understanding does not exist in free-spawning species like *Fucus*, it could still occur at the level of gametes or post-fertilization. Evidence for ‘gamete-mediated mate choice’ (GMMC) and the evolutionary significance of non-random interactions among gametes to the evolutionary origins of more definite forms of mate choice was recently reviewed (Kekäläinen and Evans 2018). On the other hand, sperm competition would be facilitated in the water column, where ejaculates from different males mix and compete for fertilization of the egg.

To test the hypothesis that the sexualization of the transcriptomes of the *Fucus* species was associated with increased sexual selection in males, we need to compare our data with transcriptomic data from closely related hermaphrodite species. For example, gene expression data from members of the two Fucales families that remained monoecious (Sargassaceae and Notheiaceae) could serve as a baseline to assess the direction of changes in expression that led to sex-bias in *F. serratus* and *F. vesiculosus* (Heesch et al. 2021).

In contrast to MBGs, female-biased genes seemed to be uniformly and highly expressed throughout the female and male body. This overall homogeneous expression pattern of FBGs became apparent when vegetative and reproductive tissue within each sex were compared (so called tissue-biased expression, as opposed to sex-biased expression where the same tissue types are compared between the two sexes). The majority of FBGs did not show tissue-biased expression in females (79% in *F. serratus* and 91% in *F. vesiculosus*), and only 20 and 22 genes showed sex-bias in vegetative tissue in *F. serratus* and *F. vesiculosus*, respectively. To summarize, sexual conflict over gene expression in *Fucus* appear to be resolved by the down-regulation of expression of pleiotropic female genes in male receptacles and by restricting the expression of MBGs to the male reproductive tissue, resulting in masculinized transcriptomes as previously reported for the giant kelp *Macrocystis* (Müller et al. 2021).

### High conservation of male-biased expression

Male-biased genes are highly conserved between the two *Fucus* species, which contrasts with the overall trends found in other species. MBGs in *Fucus* presented not only the conservation of bias, but also of expression levels (measured as Euclidean distance) which resulted in clustering of the male reproductive samples by sex rather than by species. The changes in male-biased gene regulation may have risen in the common ancestor of *F. serratus* and *F. vesiculosus* and shared ancestry could be, therefore, responsible for the observed correlation. This would further support a hypothesis that dioecy was the ancestral state in the *Fucus* genus and hermaphroditism in *F.distichus* and *F.spiralis* is a derived state. However, previous reports have shown that the targets of sex-biased expression can change over a short evolutionary time and that a small fraction of genes show parallel changes in recently diverged species (Ranz et al. 2003; Harrison et al. 2015; Huylmans et al. 2017). Similarly, studies on plants failed to find genes that were consistently sex-biased but, instead, concluded that the sex-biased gene expression was unique in each species (Scharmann et al. 2021).

Given the relatively young evolutionary age of our system, phenotypic differences accumulated between and within species may be insufficient to drive the turnover of sex-biased genes. However, this is unlikely since the number of single copy orthologs with male-biased expression (in both species) exceeded four times the number of unbiased genes with one-to-one orthologs, suggesting that the MBGs are selectively conserved. Functional analysis of male-biased genes in both species further support this assumption, as MBGs were consistently enriched in ontologies that would be important to male fertility, related to sperm production, motility and function. In contrast to MBGs, FBGs experienced higher turnover rates and had species specific expression patterns indicated by significantly increased Euclidean distance values, compared to both unbiased and male-biased genes. Taken together, if intralocus conflict is the main driver of sex-biased expression, our results suggest that the targets of this conflict are fixed in males, but not in females of *Fucus serratus* and *Fucus vesiculosus*.

### Evolution of sex-biased genes

Sex-biased genes, tend to evolve faster than unbiased genes, and this is particularly true for male-biased genes (Meiklejohn et al. 2003; Harrison et al. 2015; Lipinska et al. 2015; Darolti et al. 2018). Although male-biased genes displayed conserved expression between *Fucus* species, they presented higher rates of protein evolution compared to unbiased genes, in *Fucus serratus* and *Fucus vesiculosus*. Both, positive selection in males or relaxed selection in females may be responsible for rapid DNA sequence evolution of MBGs (Zhang et al. 2004b; Dyken and Wade 2010; Gershoni and Pietrokovski 2014; Gossmann et al. 2014; Mank 2017). Accordingly, we found 16 MBGs (30% of analyzed male-biased genes) with signatures of positive selection. Interestingly, two of these genes were associated with the sperm flagella suggesting that positive selection of male-biased genes could result from stronger sexual selection driven by *e.g.* sperm competition in *Fucus*. The fraction of MBGs under selection was, however, not significantly different to that observed for unbiased genes, indicating that adaptive evolution contributes partially, but is not the main driver of the elevated substitution rates in MBGs.

Alternatively, other aspects of genetic architecture could be contributing to the rapid evolution of male-biased genes. For example, MBGs could be less constrained by pleiotropy, because their expression is predominantly confined to male reproductive tissue, which is often associated with patterns of faster sequence evolution (Meisel 2011; Grath and Parsch 2012; Darolti et al. 2018). In line with this, female-biased genes in *Fucus* are expressed in both vegetative and reproductive tissue in male and female gametophytes, and show lower rates of synonymous to non-synonymous substitutions. Further, the rate of evolution could be determined by the genomic location of MBGs, specifically the sex-chromosome linkage. Elevated rates of coding sequence evolution on the sex chromosome relative to autosomes have been reported for several species, consistent with the theoretical prediction of fast-X o or fast-Z evolution (Kirkpatrick and Hall 2004; Mank et al. 2010; Belleghem et al. 2018). In *Fucus*, male-biased genes show high expression levels only in the male reproductive tissue, and the fast-X theory predicts that genes highly expressed in the hemizygous sex should be especially prone to fast-X evolution (Meisel et al. 2012). This interesting aspect of MBGs evolution should be revisited in the future when the genome sequences of *Fucus serratus* and *Fucus vesiculosus* become available. Finally, the set of MBGs could be enriched for young genes, which are known to evolve more rapidly (Gossmann et al. 2016). However, to assess the evolutionary age of *Fucus* sex-biased genes, additional data from closely related species is needed.

In summary, MBGs and FBGs in *Fucus* seem to follow different evolutionary path and are under different selective pressure. MBGs evolve faster at the level of the protein sequence, but their expression levels remain very conserved between *Fucus* species. In contrast, FBGs do not show accelerated rates of coding sequences evolution, but rather higher diversification of their expression levels. Because the changes in coding and changes in regulatory sequences are often decoupled, it has been suggested that they play different evolutionary roles in the evolution of morphological and physiological characters (Connallon and Knowles 2005; Wray 2007; Tirosh and Barkai 2008; Liao et al. 2009, Loehilin 2019, Martin 2013). Both types of changes (morphological or physiological) could be under selection due to reinforcement, since members of both linages (*F. serratus-F. distichus* and *F. vesiculosus–F.spiralis*) show signatures of ongoing or past hybridization, and hybrids of the dioecious *F. serratus-F. vesiculosus* are extremely rare (Coyer et al. 2002; Wallace et al. 2004; Billard et al. 2005; Coyer et al. 2007; Hoarau et al. 2015). Furthermore, studies of geographical hybrid zones of *F. serratus* and *F.distchus* (Lineage 1) show signatures of reinforcement of pre-zygotic isolation namely decreasing rates of hybridization and interspecific fertilization success with increasing duration of sympatry (Hoarau et al. 2015). Further studies are needed to characterize the genetic basis of reproductive isolation in *Fucus* as well as the connection between prezygotic barriers to fertilization and within-species sexual selection.

## MATERIALS AND METHODS

### Sampling

Four species of *Fucu*s were collected from the intertidal shoreline at Mjelle, Norway (67°24’47.3”N 14°37’49.3”E) in May 2017 (Table S1); two dioecious–*F*. *serratus and F*. *vesiculosus–and two monoecious–F. distichus* and *F*. *spiralis*. All four species were reproductively mature. To ensure availability of three replicates from each tissue type, (vegetative and reproductive tissues from males and female s for dioecious species), 10 individuals were collected from each monoecious species (*F. spiralis and F. distichus*) and 20 from each dioecious species (*F. serratus and F. vesiculosus*). Samples were transported to the laboratory in a cooler box within an hour of collection, in autoclaved seawater at 4ᵒC. The dioecious species were sexed by confirming the presence of antheridia (male) or oogonia (female) in the receptacles with a microscope. Receptacles and small segments of vegetative tissue were dissected from both monoecious and dioecious individuals and wrapped separately in aluminium foil to prevent contamination and for optimum flash freezing conditions in liquid nitrogen. All samples were stored at -80ᵒC, then freeze-dried using a VirTis Bench Top K Freeze dryer before subsequent RNA extraction.

### RNA extraction, library preparation and sequencing

Total RNA was extracted from 5mg of freeze-dried sample from every biological replicate as described in (Pearson et al. 2006). Samples were purified with the ZR-96 RNA Clean & Concentrator kit (Zymo Research, Irvine, USA) and potential PCR inhibitors were removed with the OneStep-96TM PCR Inhibitor Removal Kit (Zymo Research). RNA concentrations were quantified with the Qubit RNA Assay kit (Life Technologies, Paisley, UK) using a Qubit 2.0 Fluorometer and tested for both quantity and integrity using RNA screen tape (Agilent Technologies, Waldbronn, Germany) on the Agilent 2200 Tapestation.

Sequencing libraries were prepared from 1µg RNA using the NEBNext Ultra II Directional RNA Library Prep Kit for Illumina (New England Biolabs). Incubation time for fragmentation of total RNA was 6 mins and no size selection was performed. The libraries were sequenced twice on the Illumina NextSeq 500 (150-bp pair-end reads), using the NextSeq 500/550 High Output Kit v2.5 (300 Cycles).

### RNAseq analysis and *de novo* reference transcriptome assembly

Sequencing data were demultiplexed using the Bcl2Fastq Conversion Software (v. 2.20, Illumina). Raw sequences were adapter- and quality-trimmed with Trimmomatic (v. 0.33) (Bolger et al. 2014), followed by a standard quality check using FastQC (v. 0.11.4) (Andrews 2010), to control for aberrant read base content, length distribution, duplication and read over-representation. Prior to *de novo* transcriptome assembly, the reads were normalized to reduce redundancy of overrepresented sequences, using Trinity’s *in silico* read normalization utility (v. 2.8.5). A reference transcriptome per species was generated from the normalized reads (all replicates and conditions combined), using Trinity’s *de novo* assembly (Grabherr et al. 2011; Haas et al. 2013). Isoforms were collapsed into single gene sequences using a Trinity_gene_splice_modeler.py script from the Trinity toolkit and used for downstream analyses.

The predicted genes generated from the *de novo* assembly were then blasted against a custom bacterial/reference genomes database to identify and eliminate bacterial contamination. The longest open reading frames (ORFs) were constructed using Transdecoder (v. 5.5.0) (Haas et al. 2013), to reduce the redundant non-coding regions in the sequences. The ORFs were then blasted against an in-house heterokont database and a standard UniProt and Pfam database to keep the most likely ORFs. Transdecoder.Predict was used to predict the best coding regions with homology search results (Pfam and heterokont results) and genes without a coding region of at least 100bp were removed from the dataset. Trinity’s CD-HIT-EST (v. 4.6) (Li et al. 2001) clustered genes with predicted ORFs to further reduce the number of redundant sequences, thus generating the final reference gene sets for each species. Transcript abundances were then quantified using Kallisto (Bray et al. 2016) with 1000 bootstraps and represented as TPM (transcript per million). Genes with log_2_(TPM+1)<1 were considered not expressed.

Orthofinder (v. 2.3.3) (Emms and Kelly 2019) was used to find orthologous genes between all four *Fucus* species (Table S1). Orthofinder assigns homologous proteins (orthologs and/or paralogs) to orthogroups (OGs), so that one or more of the studied species are represented in each OG by a single-copy or multi-copy genes of common ancestry. Orthogroups with multi-copy genes allow to detect gene duplication events and provide information about conservation of genes of interest across species. They are not, however, suitable for direct gene comparisons (sequence evolution; expression levels) since it is not possible to discriminate between paralogous genes. Therefore, we used OGs with single and/or mulicopy-genes to study global patterns of conservation of sex-biased expression in the dioecious species pair; and OGs with strictly single copy genes for the evolutionary and comparative expression analyses. Orphan genes (i. e., taxonomically restricted genes) were defined as genes present in the reference transcriptome of only one species and having no BLASTp match (10−04e value cutoff) in the other *Fucus* species.

### Differential gene expression analysis

Differential gene expression within species (between sexes and tissue types) was tested with the DESeq2 package (v. 3.9 bioconductor) (Love et al. 2014). Genes with fold change FC>=2 and p_adj_ <0.05 were considered significantly differentially expressed (Benjamini-Hochberg method) and were labeled with corresponding tissue types (male, female, reproductive and vegetative). Those genes that did not show significant differential expression were labeled unbiased.

### Evolutionary analysis

Amino acid sequences of the single copy orthologous genes between *F. serratus*-*F.distichus* and *F. vesiculosus*-*F.spiralis* were aligned using MAFFT (v. 7.450) (Katoh et al. 2002) and translated back to nucleotide alignments using Pal2Nal (v. 14) (Suyama et al. 2006). The alignments were trimmed using Gblocks with a minimum block length of 20. In order to remove poorly aligned sequences that could bias the evolutionary analysis, we realigned all the fasta files with EMBOSS Water (version 6.6.0) (Madeira et al. 2019) and removed alignments with <80% similarity. The remaining high quality, gapless alignments exceeding 100 bp in length were retained for pairwise dN/dS (ω) analysis using phylogenetic analysis by maximum likelihood (PAML4, CodeML, F3x4 model of codon frequencies, runmode = − 2) (Yang 2007).

The positive selection analysis was carried out using CodeML (PAML4, F3x4 model of codon frequencies) using single copy orthologs of the four *Fucus* species (*F. serratus, F. vesiculosus, F.distichus* and *F.spiralis*) and two other brown algal species (*Ectocarpus sp.* (Cock et al. 2010) and *Sacchraina japonica (Ye et al. 2015)*). Protein alignment and curation was performed as described above. Gapless alignments longer than 100 bp containing sequences from all six species were retained for subsequent analysis. CODEML paired nested site models (M1a, M2a; M7, M8) (Yang 2007) of sequence evolution were used, and the outputs compared using the likelihood ratio test.

Euclidean distances were estimated for all single copy orthologs between *F. serratus* and *F. vesiculosus* following the approach of (Pereira et al. 2009). The following formula was used:

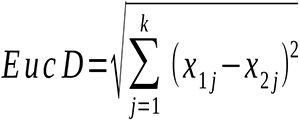

where *x_ij_* is the expression level of the gene under consideration (TPM) in species i (i.e., species 1 or species 1. 2) during stage *j* and *k* is the total number of stages (i.e., four, male and female individuals, reproductive and vegetative tissues). All statistical analysis was performed using RStudio (R version 3.6.3).

### Gene Ontology analysis

EggNOG v5.0 (Huerta-Cepas et al. 2019) was used to perform functional annotation of *F. serratus* and *F. vesiculosus* genes. We used topGO package in R (Alexa and Rahnenfuhrer 2020) to detect enrichment of specific GO terms in sex-biased genes (Fisher’s exact test with a p*-*value cutoff of 0.05).

## Acknowledgments

This work was supported by NORD University, Norway (PhD grant to WJH). We thank the CNRS-UPMC ABiMS bioinformatic platform at the Station Biologique de Roscoff, France (http://abims.sb-roscoff.fr) for providing computational resources.

Sequencing data have been deposited in the National Center for Biotechnology Information database under BioProject ID PRJNA731608.

## Supplementary data

**Table S1.** Sequencing and assembly summary.

**Table S2.** Number of sex-biased and tissue-biased genes in *F. serratus* and *F. vesiculosus; DESeq2 (FC>=2, pdj<0.05)*.

**Table S3.** Expression levels (log_2_(TPM+1) and fold change (logFC) of sex-biased and tissue-biased genes in *F. serratus* and *F. vesiculosus*.

**Table S4.** Gene orthology statistics.

**Table S5.** Orphan genes among the sex-biased genes in *F. serratus* and *F. vesiculosus*.

**Table S6**. Evolutionary rates measured as dN/dS between species pairs (*F. serratus*/*F. distichus* and *F. vesiculosus*/*F. spiralis*) for unbiased, female-biased, and male-biased genes.

**Table S7.** Gene Ontology enrichment of the sex-biased genes in *F. serratus* and *F. vesiculosus*, Fisher exact test, p<0.01.

**Table S8.** Positive selection analysis; PAML codeml with the F3X4 substitution model.

**Figure S1.**
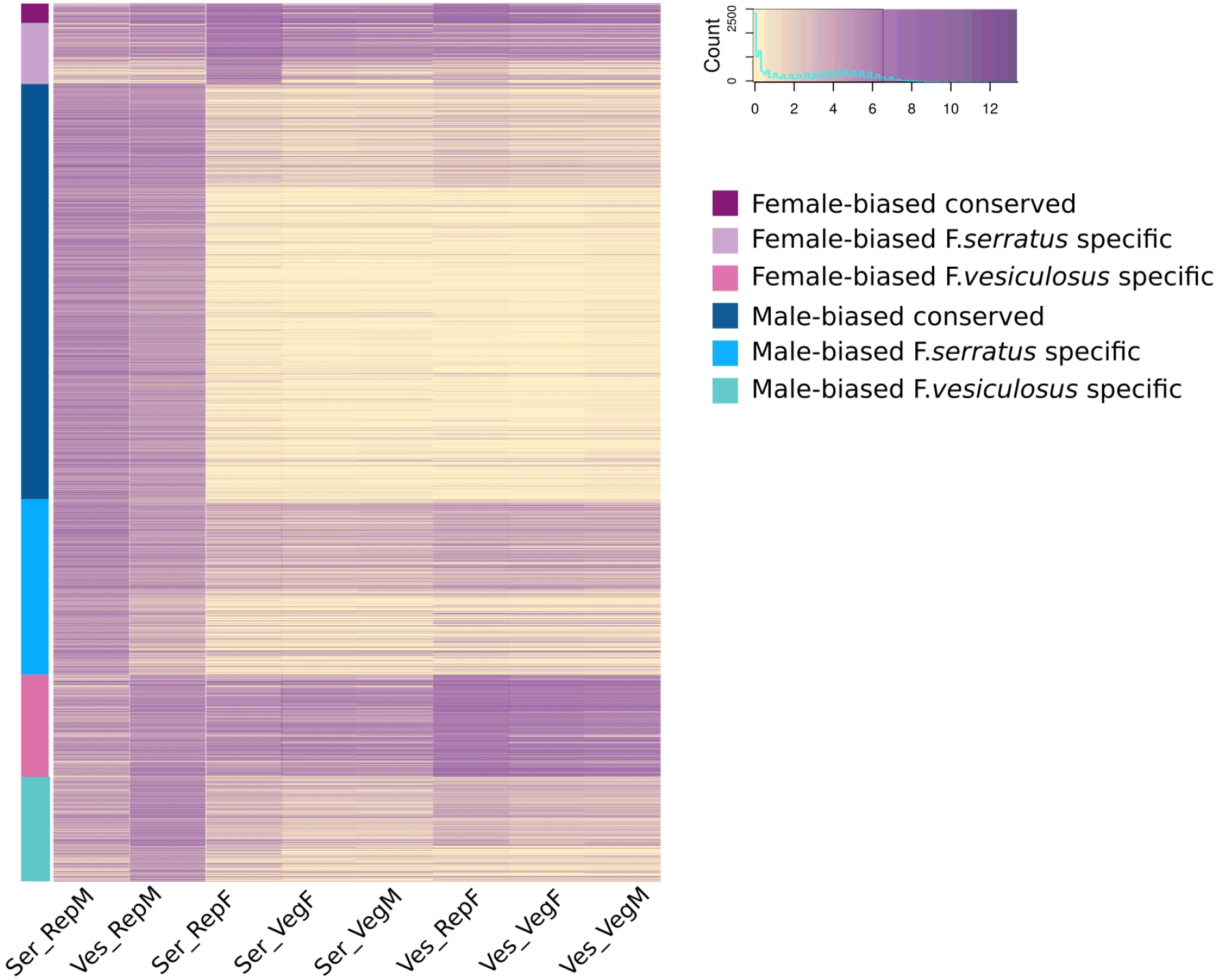
Heatmap of gene expression levels (log_2_(TPM+1)) of sex-biased genes among single copy orthologs in *F. serratus* and *F. vesiculosus* (at least one of the orthologs is sex-biased in one of the studied species). Orthologs are grouped by the conservation of sex-biased expression indicated by the colour panel on the left. Rep – Reproductive tissue, Veg – Vegetative tissue.

**Figure S2.**
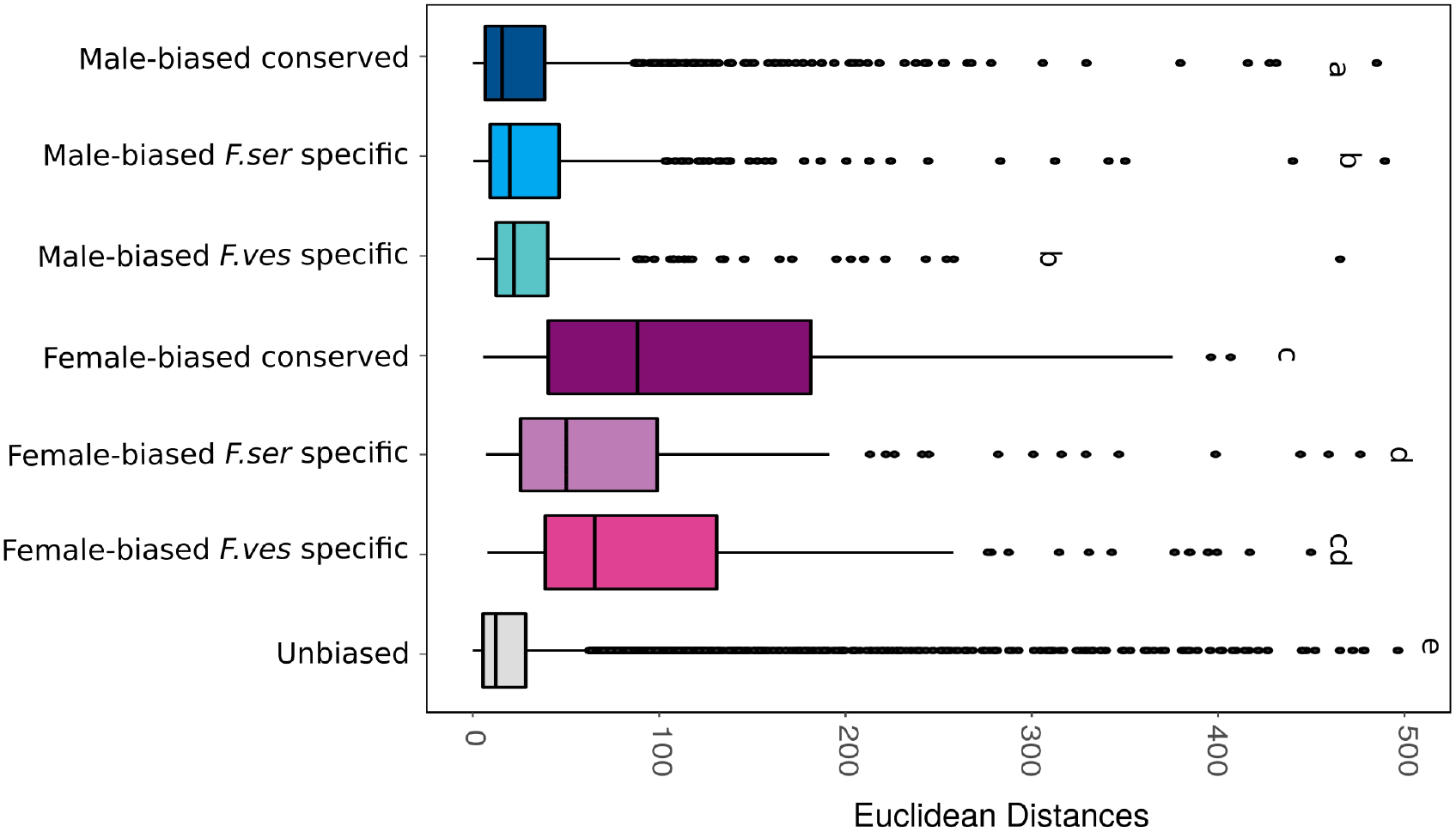
Expression divergence measured as Euclidean distances between single copy orthologous genes of *F. serratus* and *F. vesiculosus*. Sex-biased genes are divided into groups based on their conserved (orthologs show sex bias in both species) or species-specific (only one of the orthologs is sex biased in either *F. serratus* or *F. vesiculosus*) expression. Different letters above the plots indicate significant differences (pairwise Wilcoxon test; p<0.05).

